# Molecular Dynamics Simulations of Glycosaminoglycan Oligosaccharide Using Newer Force Fields

**DOI:** 10.1101/561969

**Authors:** Balaji Nagarajan, Nehru Viji Sankaranarayanan, Umesh R. Desai

## Abstract

Heparin/heparan sulfate (H/HS) are ubiquitous biopolymers that interact with many proteins to induce myriad biological functions. It is critical to understand conformational properties of H/HS in solution so as to identify their preferred protein targets. Unfortunately, the massive heterogeneity of H/HS precludes the use of solution-based experimental techniques for the thousands of sequences that occur in nature. Computational simulations offer an attractive alternative and several all-atom force fields have been developed to understand their conformational properties. Recently, CHARMM36 carrying parameters for *N*-sulfamate was developed. This work compares molecular dynamics simulations of a hexasaccharide (HS06) using two all-atom force fields – CHARMM36 and GLYCAM06. We also introduce two new straightforward parameters, including end-to-end distance and minimum volume enclosing ellipsoid, to understand the conformational behavior of HS06. In addition, we analyzed inter- and intra-molecular hydrogen bonds and intermediate water bridges formed for HS06 using both force fields. Overall, CHARMM36 and GLYCAM06 gave comparable results, despite few, small differences. The MD simulations show that HS06 samples a range of conformations in solution with more than one nearly equivalent global minima, which contrasts with the assumed single conformation conclusion derived on the basis of 1HPN structure. A key reason for the stability of multiple low energy conformations was the contribution of intermediate water bridges, which is usually not evaluated in most MD studies of H/HS.

## Introduction

Heparin/heparan sulfate (H/HS) are linear polysaccharides that bind many proteins and induce important biological functions such as growth, differentiation, inflammation, adhesion, and many more ^1–5^. The affinity and specificity of these highly anionic polysaccharides arise from their differential local sulfation level, especially in constituent di-, tetra- and hexa-saccharide blocks consisting of iduronic acid (IdoA)/glucuronic acid (GlcA) and glucosamine (GlcNAc) (Figure 1) ^6, 7^. The number of variations possible for these oligosaccharides is very high, which makes it extremely difficult to study individual sequences in solution ^8^.

**Figure 1.**
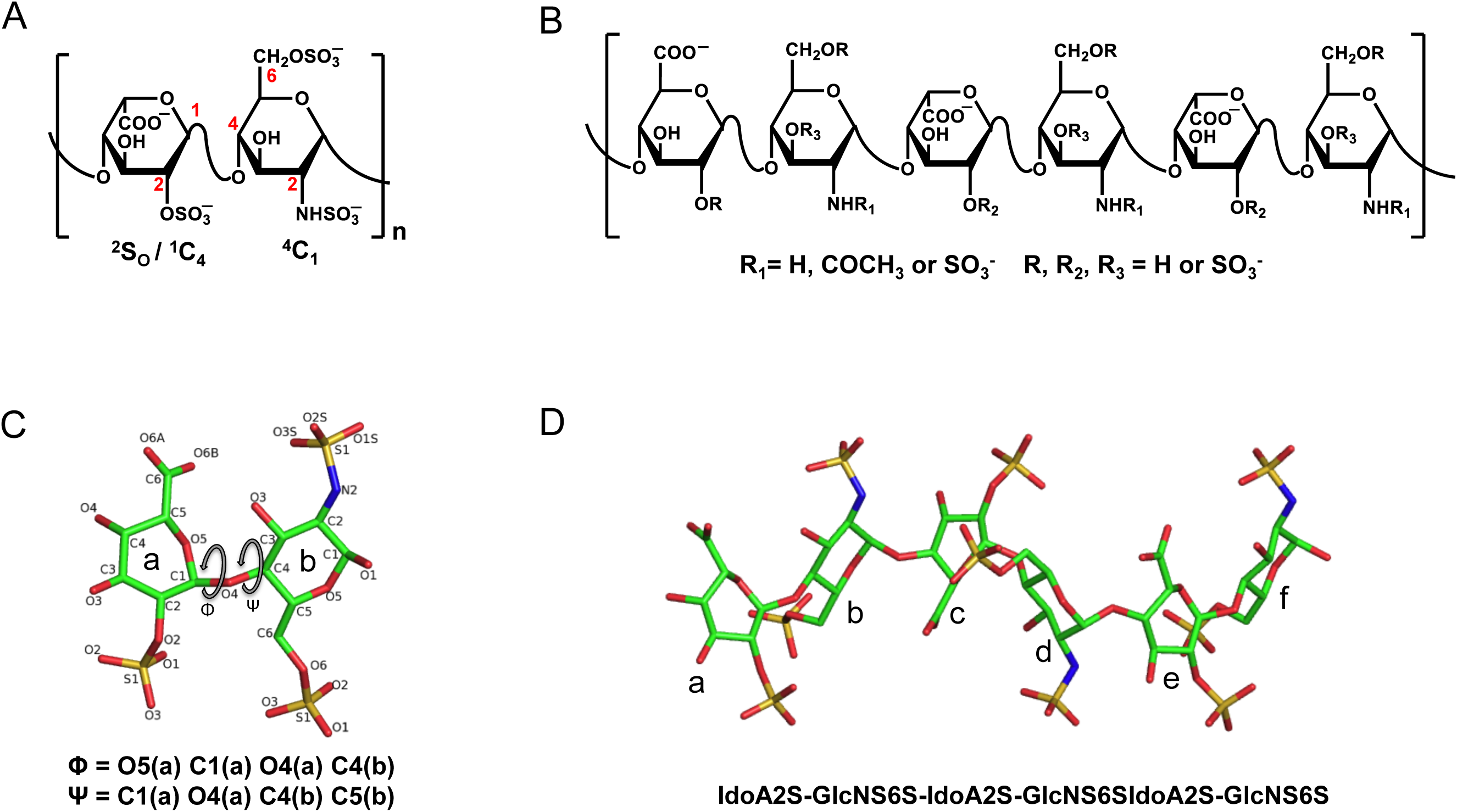
A) A two-dimensional representation of heparin polymer showing a common building block IdoA2S–GlcNS6S. IdoA could be in ^2^S_O_ or ^1^C_4_ pucker, while GlcNAc is in ^4^C_1_ pucker. Positions are marked in red. B) Heparin/heparan sulfate are heterogeneous entities with variable sulfation and chain length. Shown is a hexasaccharide chain (HS06) with variable substitutions. C) A disaccharide unit of IdoA2S(^2^S_O_)–GlcNS6S(^4^C_1_) shown with all non-hydrogen atoms and torsional angles Ф and Ψ. D) The structure of HS06 deposited in the PDB (ID: 1HPN) with residues *a*, *c*, *e* being IdoA2S and *b*, *d*, *f* being GlcNS6S (hydrogens are not shown for clarity).

To overcome the difficulty of studying individual sequences in solution, in silico techniques such as molecular modeling and molecular dynamics (MD) are very attractive. Both techniques are relatively inexpensive and can help quickly screen numerous sequences for desired properties, e.g., identifying their propensity to recognize a target binding site ^9, 10^. When using MD, it is also possible to study conformational transitions at microsecond time scale, which are difficult to study using other techniques ^11–13^. More importantly, MD can help deduce atomistic information that explains unique interaction of drugs with biomacromolecules. With regard to H/HS, MD has been used to study the affinity of conformations for proteins ^14–17^, which implies that MD studies could become key to designing H/HS-based therapeutic agents.

Yet, appropriate tools have not been developed to make MD easy to implement for a large library of H/HS sequences. Additionally, H/HS building blocks appear to deceptively simple, but are substantially flexible ^18, 19^, which makes MD simulations challenging. In fact, a number of studies on H/HS have attempted to elucidate conformational flexibility of IdoA by comparing computational results with solution ^1^H NMR results ^20–24^. This has offered a strategy to fine-tune parameters for sulfated monosaccharide residues. As a result, many computational studies of H/HS conformational profiles in water and in the protein bound form have been presented ^10, 12–5, 25–31^. The majority of these studies have utilized either GLYCAM, CHARMM or GROMOS force fields, which are the primary force fields available to study carbohydrates ^32–34^.

In this work, we present detailed procedures to perform an all-atom MD simulation of an H/HS oligosaccharide using the two recent force fields, GLYCAM06 and CHARMM36. H/HS hexasaccharide (HS06), derived using the 1HPN conformation ^22^ in which all IdoA2S residues are in the ^2^S_O_ form has been used as a test oligosaccharide. We have evaluated the similarities and differences in torsional angle variations, inter and intra-molecular hydrogen bonds, and intermediate water bridges between the two force fields. In this paper, we also introduce two new straightforward parameters to understand the conformational behavior of HS06 at a molecular level. We find the both force fields predict that HS06 samples a range of conformations in solution with more than one nearly equivalent global minima, which is at variance from the conclusion derived from the single conformation shown by the 1HPN structure. Both force fields gave comparable results except for few, small interesting differences. We expect these results to explain the plasticity of interactions in terms of GAG recognition by proteins ^17, 35^, while also greatly aiding discovery of GAGs as modulators of protein function ^9^.

## Materials and Methods

The primary all-atom force fields available for conformational simulations of GAGs include GLYCAM, CHARMM and GROMACS ^32, 34, 36^. We selected GLYCAM06 and CHARMM36 for this study. Recent additions to the CHARMM force field include parameters for *N*-sulfamate ^37^, which were implemented in this study. Likewise, recent additions to the GLYCAM force field include parameters for Δ^4, 5^ unsaturated uronate (ΔUA) ^38^. MD simulations were performed using NAMD program 2.9 employing CHARMM36 force field ^39^, whereas simulations employing GLYCAM06 force field were performed using AMBER simulation package ^40^.

### GLYCAM06/AMBER Preparation Step

MD studies of HS06 in explicit water were performed using AMBER14 ^40^ with GLYCAM06 force field parameters ^32^. The initial HS06 structure was the experimental NMR structure reported in the protein data bank (PDB, ID: 1HPN) ^22^. This structure has a repeating sequence of (IdoA2S-GlcNS6S)_3_ with IdoA2S in skew boat ^2^S_O_ conformations (Figure 1). The building blocks for this sequence were identified from the library in GLYCAM and appropriate residue/atom labels were re-named to conform to the parameter files (refer to http://glycam.org/docs/forcefield/glycam-naming-2/). The HS06 structure was loaded in xleap and analyzed for the glycosidic linkage connectivity as well as preferred atom- and residue-types using the ‘edit’ function in xleap (see http://ambermd.org/tutorials/). This was followed by ensuring that the overall charge of HS06 is −12 (use the ‘charge’ command) and adding 12 Na^+^ counter ions to ensure an overall net charge of zero. The system was then solvated using TIP3P water molecules using the ‘solvatebox’ function, wherein the distance between any atom of HS06 and the nearest box wall was 10 Å ^40^. The initial parameter and topology for the system were saved for future steps. These details are presented as a flow chart in Supporting Materials Figure S1A.

### GLYCAM06/AMBER Minimization Step

Following set up of initial parameters and co-ordinate files, a two-step minimization procedure was implemented. In the first step, the solute and the Na^+^ ions were restrained using a harmonic potential of 100 kcal/(mol Å^2^). The water molecules were relaxed using 500 cycles of steepest descent and 1500 cycles of conjugate gradient method. In the second step, the whole system was relaxed to 2500 cycles of conjugate gradient minimization. The system was then brought to a constant temperature of 300 K using the temperature coupling with a time constant of 2 fs followed by achievement of a constant pressure of 1 atm. Finally, the system was equilibrated to NPT. These phases were implemented for a total time period of 1 ns with 2 fs of integration time step.

### GLYCAM06/AMBER MD Run

The MD simulations were then initiated for a total time period of 20 ns (integration time step = 1 fs) during which the ensemble coordinates were collected at every 1 ps. During this simulation, the periodic boundary condition, particle-mesh Ewald method and a non-bonded cut-off of 10 Å were employed. The covalent bonds involving hydrogen atoms were constrained using the SHAKE algorithm throughout the simulation^40^. A weak torsional restrain was applied to keep the puckering of IdoA2S in ^2^S_O_ conformation throughout the simulation ^25^.

### CHARMM36/NAMD Preparation Step

The initial HS06 structure used in these experiments was the same as for GLYCAM06/AMBER studies (Figure 1), except that residue and atom labeling were changed to match with CHARMM36 program (see http://mackerell.umaryland.edu/charmm_ff.shtml). The initial structure of HS06 was loaded in VMD using the ‘Tk’ console with par_all36_carb.prm topology file ^41^. The patches for sulfation were introduced and the structure validated for absence of any errors. The total charge on the system was then ascertained using the protein structure file (psf), the coordinate file (pdb) and the ‘get_total_charge’ script from VMD archive library (http://www.ks.uiuc.edu/Research/vmd/script_library/scripts/total_charge), which gave an integral value of −12, as expected. HS06 was then solvated using the ‘solvate’ package using TIP3P box measuring at least 10 Å from an HS06 atom to the nearest box wall. Finally, 12 Na^+^ cations were added using the ‘autoionize’ function to neutralize the system. The initial protein structure (psf) and the coordinate (pdb) files of HS06 in solvent were saved for further procedures (see Supporting Figure S1B).

### CHARMM36/NAMD Minimization Step

CHARMM36/NAMD minimization was performed in two steps, as described for minimization in GLYCAM06/AMBER (above). Briefly, each step involved a total of 2000 conjugate gradient steps. The system was equilibrated in three steps to constant temperature 300 K and pressure of 1 atm in an NPT ensemble for a total of 1 ns, with integration time step being 2 fs.

### CHARMM36/NAMD MD Run

Following equilibration, an unrestrained MD simulation was performed for 20 ns and the MD trajectory stored for every 1 ps with 1 fs integration time step. A weak torsional restrain was applied to maintain the ^2^S_O_ pucker for IdoA2S residue throughout the simulation ^25^. Periodic boundary condition, particle-mesh Ewald method and a non-bonded cut off 10 Å with a switch distance of 8.5 Å were employed in MD runs. The covalent bonds involving hydrogen atoms were constrained using the SHAKE algorithm throughout the simulation.

### Analysis of MD data

Standard protocols from AMBER and NAMD ^39, 40^ were used for analyzing MD runs. We analyzed the final 2 ns of data to present the results in a concise manner. Conformational exploration of HS06 was studied using two recently developed tools called end-to-end distance (EED) between (α-*O*4 and Φ-*O*1, see Figure 2A) and minimal volume enclosing ellipsoid (MVEE) (see Figure 2C). In-house scripts were developed for automated extraction of EED and MVEE from the conformers generated in the MD runs^42^. We also analyzed backbone torsional angles and inter- and intra-molecular hydrogen bonds for every conformer that arose during MD run for both the force fields. In addition, we analyzed inter- and intra-molecular bridged water molecules identified in MD simulations to reflect upon the role of solvent in stability particular conformers of HS06. Finally, we analyzed the potential energy landscape of HS06 across the entire MD trajectory using parameters EED and MVEE. Programs ‘VMD’ and ‘cpptraj’ were used for trajectory analysis.

**Figure 2.**
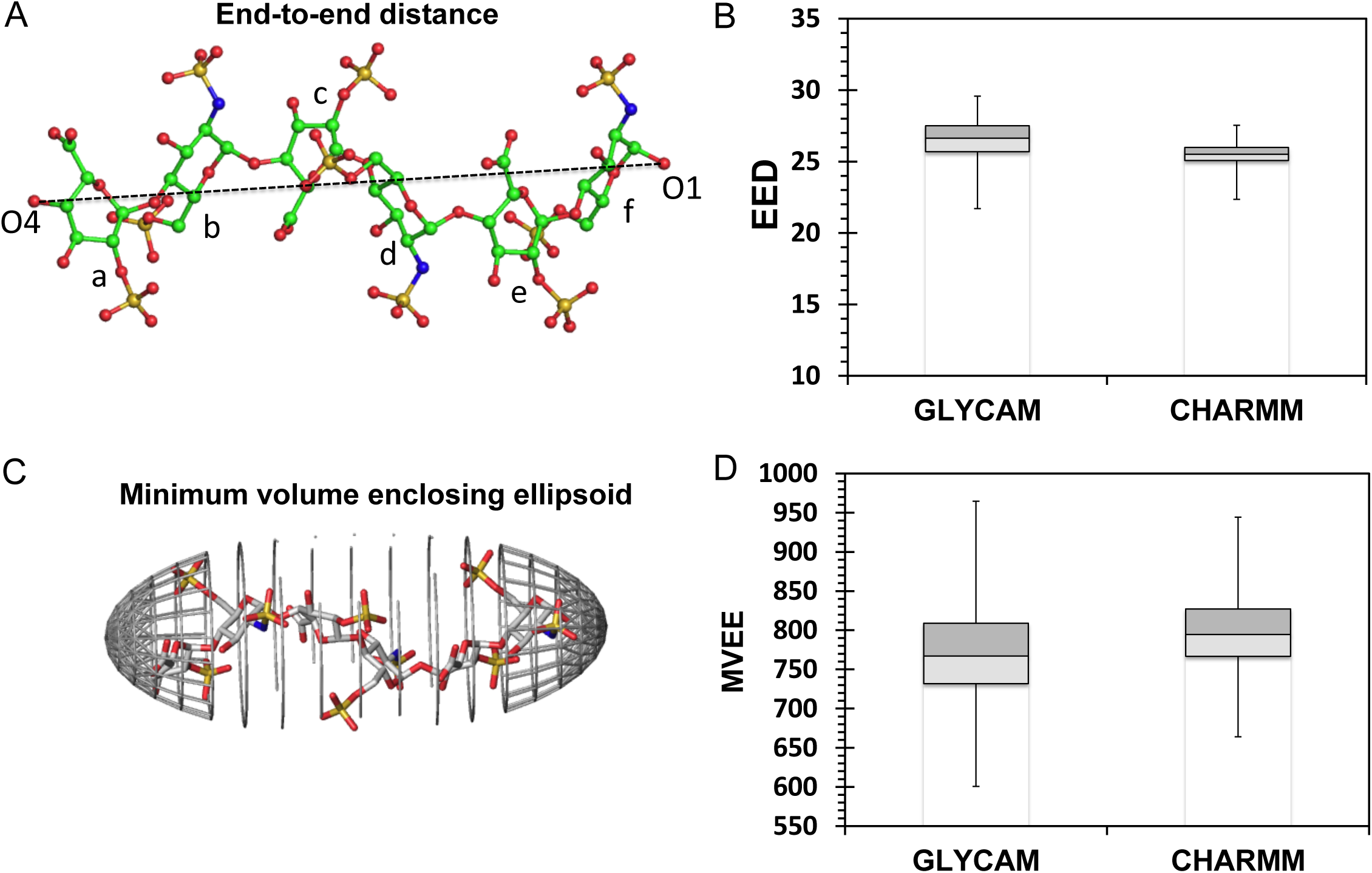
A) Representation of the end-to-end distance (EED) between terminal oxygen atoms (a@O4 and f@O1) of HS06. B) Box plot of EED (in Å) from MD generated conformations using GLYCAM06 and CHARMM36 force fields. C) Representation of the minimum volume enclosing ellipsoid (MVEE) for the coordinate space of HS06. The saccharide is shown in stick form and enclosing ellipsoid grids are shown in grey color lines (for clarity, not all the lines are shown). D) Box plot of MVEE (in Å^3^) from MD generated conformations using GLYCAM06 and CHARMM36 force fields.

## Results and Discussion

### Parameters to Assess Overall Conformation

We employed two simple parameters, end-to-end distance (EED, Figure 2A) and minimum volume enclosing ellipsoid (MVEE, Figure 2B), to help understand conformational behavior of GAG sequences, e.g., HS06, as the MD trajectory proceeds. EED approximates the linear length of a sequence and primarily reflects on the bendability, or lack thereof, of the sequence. H/HS chains typically exhibit an overall helical secondary structure, which can carry kinks and bends, if appropriate substitution pattern is present. Although the ^4^C_1_ and ^1^C_4_ forms of GlcA/GlcN and IdoA residues, respectively, introduce a more rod-like helical structure, the ^2^S_O_ form of IdoA induces significant probability of kinks and bends ^43–46^. In fact, IdoA is thought to be the key residue that enables attainment of a wider conformational space. Thus, we propose EED as a quick measure of the ‘bendability’ of a GAG sequence. For example, a rigid rod-like form will display an average EED of 100%, whereas the EED of a hairpin bend form would correspond to a ∼20% or less.

Another parameter is MVEE, which corresponds to how voluminous is the given sequence when tumbling in solution. Alternatively, MVEE conveys information on whether the sequence exhibits an average of tubular, oval or spheroidal form across the MD trajectory. For example, HS06 in a rigid rod-like form would have MVEE of ∼674 Å^3^; in contrast, a hairpin shaped HS06, which is possible only under in silico conditions, would display a MVEE of ~643 Å^3^ (see supplementary information Figure S2 and Table S1).

### CHARMM36 is very similar to GLYCAM06 with some interesting differences

We evaluated conformational space explored by HS06 through calculation of EED and MVEE for every conformer sampled in MD simulations using CHARMM36 and GLYCAM06 force fields (see Figure S3). Figures 2C and 2D show the distribution of EED and MVEE, respectively, across the trajectory for the two force fields. For both parameters, the box plots were fairly similar. The average EED using GLYCAM06 (26.5 Å) was different from that obtained using CHARMM36 (25.5 Å) by only 1 Å. Both of these compare favorably with that measured using NMR (26.4 Å) (Table S1). Likewise, the difference in average MVEE for the two force fields was only 23.9 Å^3^ (Table S1), which arose from rather similar average values of the three radii (r1, r2, r3) defining the ellipsoid. The MD derived values of the three radii are also very similar to those calculated for the NMR-derived HS06 structure (Table S1). Further, the distribution of conformers was fairly symmetrical from the two medians (EED and MVEE) for both force fields suggesting good correspondence between the two force fields as well as with biophysical measurements. The overall statistics showed that HS06 sampled ~5% bent conformers with EED less than 24 Å. Alternatively, 90% of conformers were either rod-type or rod-like in shape, which corresponded to EED of 25 to 28 Å. Throughout the simulations, HS06 was found to sample multiple conformers with the majority being similar to the solution NMR structure (PDB ID:1HPN) (see Figure S4).

Yet, there were some interesting differences. 1) The conformations sampled using CHARMM36 were found to fit well within the range of GLYCAM. Alternatively, the range of conformations sampled using CHARM36 force field was slightly lower than that using GLYCAM06. 2) The interquartile range for CHARMM36 was also less than that for GLYCAM06. This implies that CHARMM36 predicts a slightly more rigid HS06 structure in comparison to that predicted by GLYCAM06. The apparent reason for this difference is described below in the section on inter-molecular hydrogen bonding interactions.

### CHARMM36- and GLYCAM06-derived torsions are very similar

Each conformer in HS06 has a pair of phi and psi values for each of its interglycosidic linkages, i.e., three Φ and three Ψ values for both IdoA2S-GlcNS6S and GlcNS6S-IdoA2S. The calculated Φ and Ψ for all conformers along the MD trajectories are shown in Supporting Information (Figure S5 and S6). Overall, the torsion angles are in good agreement between CHARMM36 and GLYCAM06; a difference of only 2 to 30 degrees was noticed across the entire MD run, which is relatively small considering the dynamic motion of the HS06 sequence. In fact, the average values of corresponding Φ and Ψ for both inter-glycosidic linkages were −74.99° (−67.73°) and −113.18° (−115.99°) (IdoA2S-GlcNS6S) and 85.33° (65.11°) and −140.41° (−131.98°) (GlcNS6S-IdoA2S), respectively, which implies that both force fields predict nearly identical conformations across the inter-glycosidic bonds. These values correlate well with earlier results reported in the literature ^25^ and are also within the acceptable range based on the solution NMR structure ^22, 47^.

### Intra-molecular hydrogen bonds predicted by the two force fields show some difference

The stability of overall HS06 conformation is not only dependent on the glycosidic torsional space but also dependent on the formation of intra-molecular hydrogen bonds (H-bonds) within and between neighboring residues ^27, 48, 49^. The intra-molecular H-bonds for the MD trajectory were calculated using the cpptraj tool available from Amber tools 15. Herein, a H-bond is defined based upon a distance of 3.5 Å between two electronegative atoms and a deviation within ±60° from the ideal linear arrangement^50^. Figures 3A and 3B show the occurrence of the number of H-bonds across each frame of MD trajectory for GLYCAM06 and CHARMM36, respectively. The average H-bonds for GLYCAM06 were 4.3, while CHARMM36 gave an average of 3.1; the difference of 1 intra-molecular H-bond between the two force fields is not high but worth noting. This difference is also maintained if we calculate the number of H-bonds that persist over 90% of MD trajectory suggesting a measurable difference between the two force fields. However, significant H-bonds formed between residues were found to be identical across both force fields (Figure 3C and 3D) suggesting excellent correspondence between CHARMM36 and GLYCAM06. It is important to note that these specific H-bonds are identical to those reported earlier (IdoA2S O5 ▪ ▪ ▪ ▪ GlcNS6S@H3O and IdoA2S O61 ▪ ▪ ▪ ▪ GlcNS6S@H3O) ^27, 48^. Figure 3D shows the percentage occurrence of three significant H-bonds bonds and, these span equally in both the force fields.

**Figure 3.**
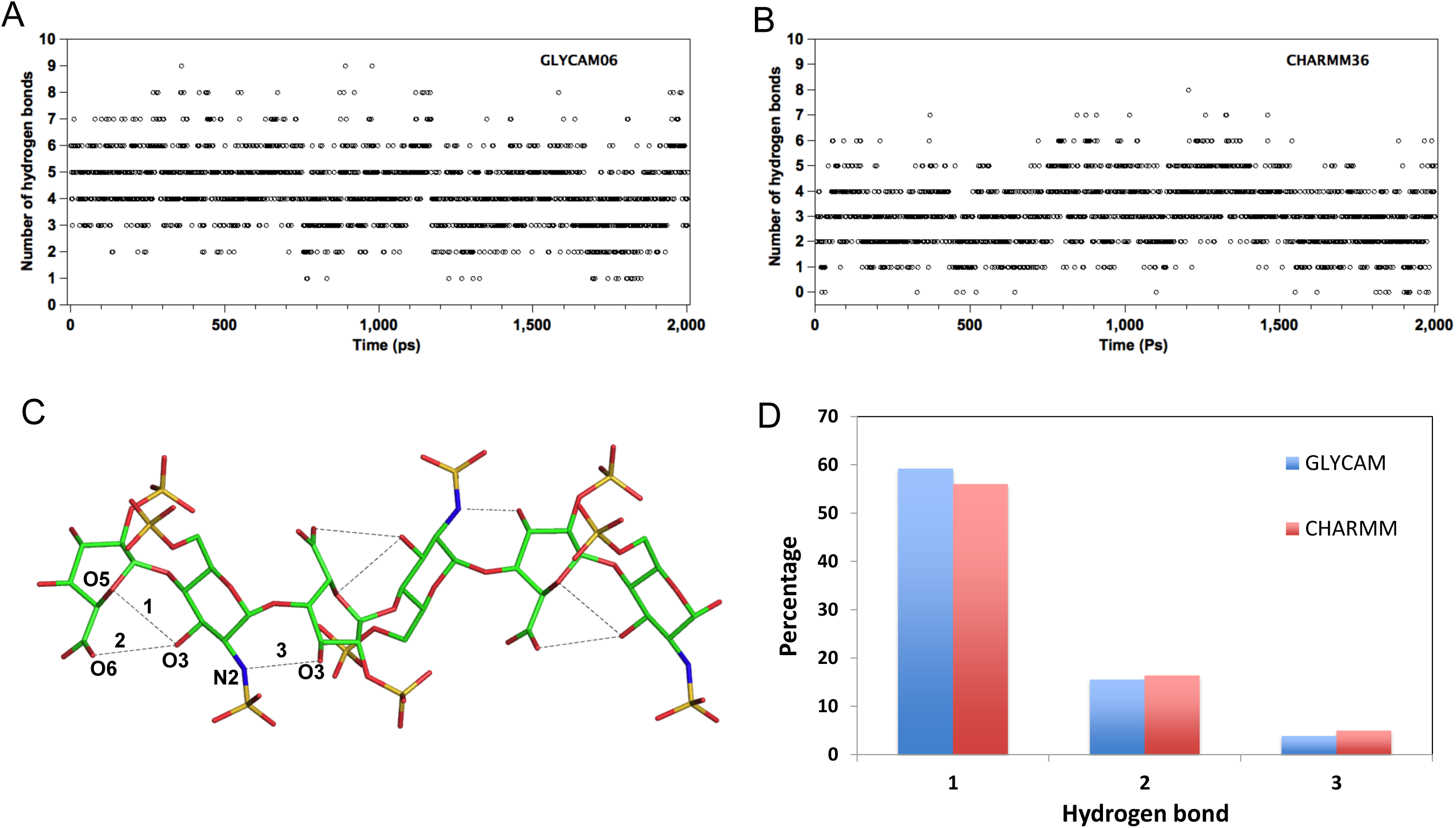
The total number of hydrogen bond between all possible donor and acceptor atoms of HS06 using A) GLYCAM generated conformations and B) CHARMM generated conformations. C) The intra-molecular hydrogen bond interactions in HS06 displayed by the two force fields. Hydrogen bonds are shown in dotted lines between the respective donor and acceptor atoms. D) The percentage occupancy of the three significant hydrogen bonds including 1) IdoA2S@O5•••GlcNS6S@H3O•••GlcNS6S@O3; 2) IdoA2S@O6•••GlcNS6S@H3O•••GlcNS6S@O3; 3) GlcNS6S@N2•••IdoA2S@H3O•••IdoA2S@O3.

### Inter-molecular hydrogen bonds predicted by the two force fields show some difference

The presence of multiple sulfate groups on a H/HS sequence introduces strong interactions with multiple water molecules ^25, 27, 51, 52^. In fact, the local conformation and flexibility (or rigidity) of a H/HS chain also depend on inter-molecular H-bonds with solvent molecules. Earlier, the number of water molecules bound to a chain was found to vary with IdoA pucker ^53^. Studies have also shown radial distribution of water around the polar groups and have confirmed that the skew boat form of IdoA is more solvated than the chair form ^25, 27^.

Figures 4A and 4B show representative frames of HS06 structure from both simulations, GLYCAM06 and CHARMM36, respectively. The snapshots display a rich bed of water molecules around HS06 involved in the formation of a large number of inter-molecular H-bonds. We calculated the number of water molecules interacting with HS06 across the two MD trajectories using cpptraj. Both force fields show an equivalent number of water molecules around the polar groups of HS06. As expected, water molecules stabilize HS06 conformation by bridging two polar groups in two different ways: a) a single water molecule forms a bridge between a donor-acceptor pair of HS06 and b) two or more water molecules form a network between donor and acceptor pair of HS06. Both these geometries are shown in Figures 4C and 4D.

**Figure 4.**
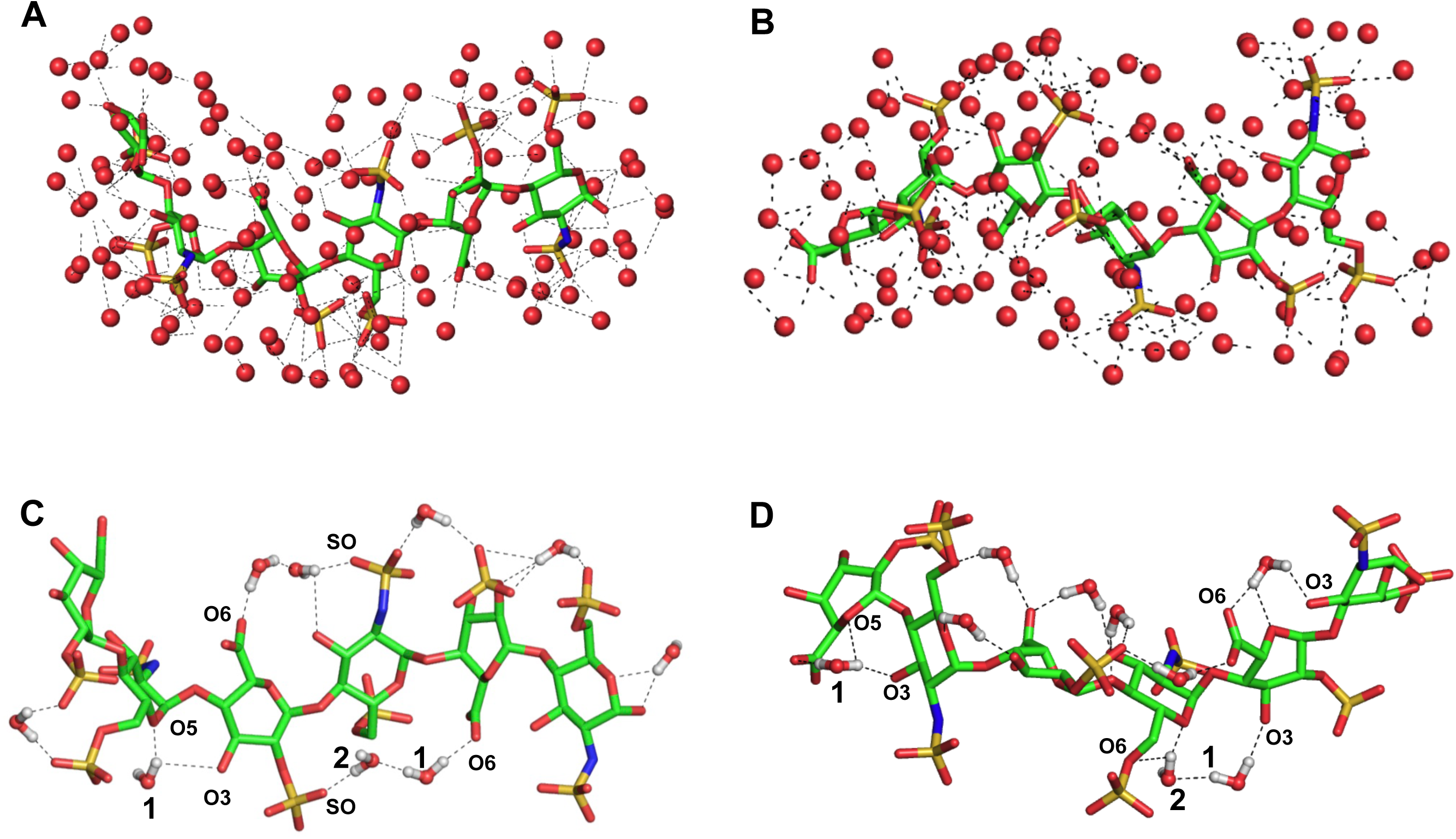
Hydration shell around HS06. The inter-molecular hydrogen bond interactions by water molecules to HS06 atoms are shown as dotted lines. HS06 is shown in stick representation and water molecules are shown as red spheres without hydrogen atoms. The conformations generated by A) GLYCAM force field and B) CHARMM force field are shown. Bridging water molecules between the polar atoms of neighboring molecules. 1 and 2 represents the water network in-between polar atoms. C) shows bridging water molecules observed from GLYCAM generated conformations, while D) shows bridging water molecules observed from CHARMM generated conformation.

The number of bridging water molecules in both simulations varied from 1 to 14 (see Figure S7). The bridging water molecules could play a critical role in protein binding in the form of imparting desolvation energy as well as stabilizing native state ^54^. It is important to note that both GLYCAM06 and CHARMM36 identified a rather identical set of bridging waters belonging to the two categories described above (see Figures 4C and 4D). Analysis of the number of bridging water molecules across MD runs shows an average of 6.09 and 4.65 for GLYCAM06 and CHARMM36, respectively. Once again, GLYCAM06 appears to identify more inter-molecular H-bonds in comparison to CHARMM36.

The difference in inter-molecular H-bonding interactions may be the reason for the conformational flexibility, or rigidity, observed for HS06 observed between the two force fields. HS06 exhibits a more rigid form according to CHARMM36, whereas it appears to sample wider range of forms according to GLYCAM06. It would be important to recognize and appropriately apply these differences in future applications of the force fields for GAG-based drug discovery.

### Energy landscape

We explored the potential energy landscape (PEL) of HS06 in three-dimensional space (3D) for the two force fields using the two parameters of our interest, EED and MVEE, rather than the typical analysis used for protein folding involving two principal coordinates ^55–57^. Figure 5 shows the two PELs for the HS06 computed using the two force fields, GLYCAM06 and CHARMM36. Both force fields exhibit multiple minima resident on a rugged landscape. The ruggedness of PEL appears to arise from fluctuations in glycosidic torsions and dynamism of multiple inter- and intra-molecular H-bond.

**Figure 5.**
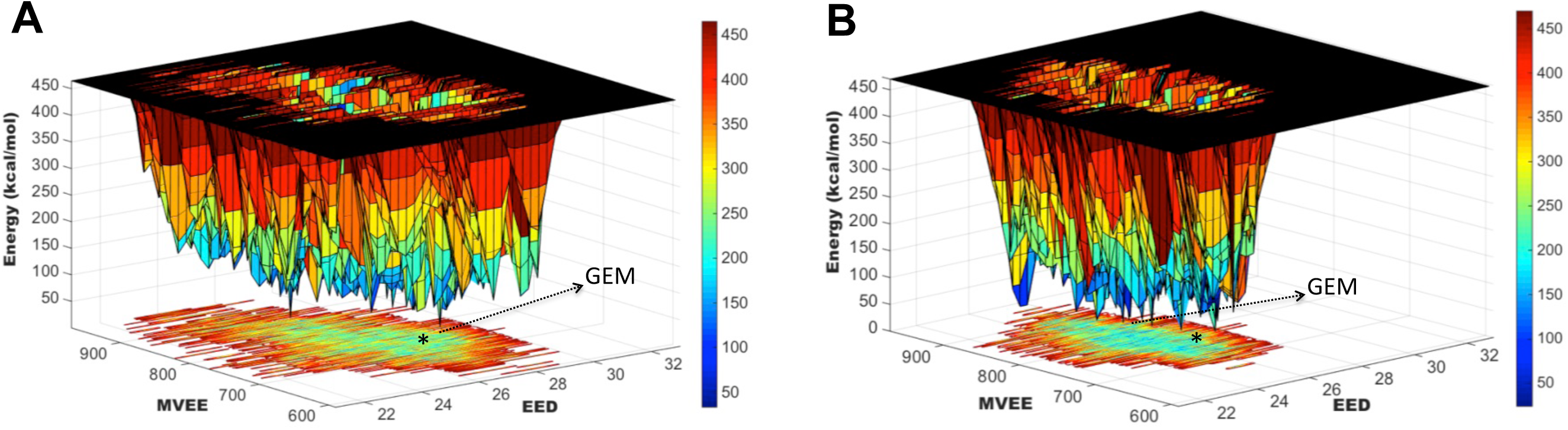
The three-dimensional (3D) potential energy landscape of the conformational sampling from the two force fields: (A) GLYCAM and B) CHARMM, using EED (in Å) as X-axis and MVEE (in Å^3^) as Y-axis. The potential energy (kcal/mol) is represented in color bars. Observations from the native 1HPN are marked as * and the global minimum energy (GEM) is shown as arrow.

Comparison of the PELs shows that GLYCAM06 invokes marginally higher conformational entropy (Figures 5C and 5D). Both force fields display a unique global energy minimum (GEM) conformer. Whereas GLYCAM06 presents it as a conformer with EED and MVEE values of 25.3 Å and 829.7 Å^3^, respectively, CHARMM36 presents it as the 24.1 Å, 804.9 Å^3^ conformer (Figure 5). Further analysis of the PELs shows that the landscape around the GEMs is rather steep and canyon-like. However, the energy difference between the ridges and the valley is not high, which implies that several local minima in the vicinity of the two GEMs could be sampled by HS06 at room temperature.

The obvious major significance of such as mobile conformational equilibrium is in the phenomenon of protein recognition. The affinity of a protein that binds to HS06 will intricately depend on the nature of this conformational equilibrium, which determines the population of the particular form of HS06 recognized. By the same token, cross-reactivity of HS06 for other proteins would also depend on this equilibrium. This implies that drug design researchers would have to factor this conformational equilibrium in their molecular modeling studies.

## Conclusions

GAGs, present in extracellular matrix, on cell surface and within cells, are functionally relevant for biological responses. For example, heparan sulfate (HS) brings about protein–protein interactions of importance to growth and differentiation ^58, 59^. Traditionally, it has always been difficult to deduce the bases of biologic signals are mediated by GAGs. We project that the approach of understanding dynamical conformational properties of GAGs would help elucidate mechanistic details on GAG recognition and modulation of proteins. For example, our earlier work on HS06 recognition of fibroblast growth factor and its receptor ^42^ yielded significant foundational understanding into how cancer stem cell growth could be modulated^60^.

The dynamical properties of individual GAGs are challenging to study using traditional biophysical tools because of structural and conformational heterogeneity of samples. To truly develop understanding on how natural GAGs, carrying an ensemble of sequences, may behave in biological fluids, we have taken the first step of analyzing MD simulations of HS06 using two newer force fields, GLYCAM06 and CHARMM36. Both force fields yielded rather similar conformational dynamism, which appeared to be comparable to earlier results from NMR solution experiments. More specifically, both gave essentially equivalent overall conformation with similar inter- and intra-molecular interactions. Yet, the simulations showed that HS06 exhibits dynamism in solution with more than one nearly equivalent global minima, which contrasts with the assumed single conformation conclusion derived on the basis of 1HPN structure.

Of particular importance is that we utilized two new parameters EED and MVEE to understand the conformational dynamism. Rather than the non-physical principal components typically used in analysis of energy landscapes, we used EED and MVEE, which can directly correlate with the conformational properties of oligosaccharides. These plots presented HS06’s local minima at preferred positions only. We expect that EED and MVEE will significantly help with the design of GAG sequences for targeting proteins.

We also project that application and analysis of EED and MVEE to libraries of GAG sequences will help correlate inter- and intra-molecular interactions, which could explain why some sequences are ‘specific’ and whereas others are ‘plastic’ in naturally occurring GAGs. Of special note is the common occurrence of a specific intra-molecular hydrogen bond arising from the 3OH group of IdoA2S, which was not affected much by the solvent molecules, resulting in a drive to ^2^S_O_ pucker ^27^. Likewise, application of these tools would perhaps explain the structure-dependent kinks and bends in HS oligosaccharides that appear to play a major role in recognition of proteins^43–46^.

## Supporting information

Supplementary Informations

## Acknowledgments

We thank Prof. Glen Kellogg and Prof. Neel Scarsdale of VCU for computational support. We also thank the availability of research resources from National Center for Research Resources (S10 RR027411) to VCU. This work was supported in part by grants from the NIH including HL107152, HL090586 and HL128639. CHiPC VCU.

