## Supplementary Informations for "Molecular Dynamics Simulations of Glycosaminoglycan Oligosaccharide Using Newer Force Fields"

for


| # | **Table of Contents** | Pg. |
|---|---|---|
| 1 | Figure S1. Steps to generate initial starting structure (GLYCAM06 and CHARMM36) | 2 |
| 2 | Figure S2. HS06 rod and hairpin bent structure | 3 |
| 3 | Table S1. Average value of end-to-end distance EED and minimum volume enclosing ellipsoid MVEE from GLYCAM06 and CHARMM36 [illegible] | 4 |
| 4 | Figure S3. End-to-end distance EED and minimum volume enclosing ellipsoid MVEE from GLYCAM06 and CHARMM36 force field simulations [illegible] | 5 |
| 5 | Figure S4. Representative conformers shown from different bins of EED | 6 |
| 6 | Figure S5. Phi psi from IdoA2S-GlcNA6S glycosidic linkage for last 2ns from GLYCAM06 and CHARMM36 force field simulations | 7 |
| 7 | Figure S6. Phi psi from GlcNA6S-IdoA2S glycosidic linkage for last 2ns from GLYCAM06 and CHARMM36 force field simulations | 8 |
| 8 | Figure S7. The number of bridging water molecule interactions | 9 |

**A**

Initial 3D structure
X-ray/NMR/Modeling

Understanding building blocks converting residue and atom labeling to GLYCAM06

Calling GLYCAM06 leaprc in xleap of AMBER14/16

Load structure (structure of interest)

Check the structure by edit command in xleap

Check for linkage and total integer charge

Add enough counter ion to charge neutralize eg. $Na^+$

Solvate all in TIP3PBOX

Save initial parameter and topology

**B**

Initial 3D structure
X-ray/NMR/Modeling

Understanding building blocks converting residue and atom labeling to CHARMM36

Open VMD Tk Console load topology file par_all36_carb.prm

Load structure (structure of interest)

Check the structure in VMD

Check for linkage and total integer charge

Solvate all in TIP3PBOX

Autoionize with counter ion to charge neutralize eg. $Na^+$

Save initial protein structure file and coordinates.

Figure S1. Steps to generate the initial starting structure for the molecular dynamics simulation A) using GLYCAM06 force field in AMBER and B) CHARMM36 force field parameter using VMD/NAMD.

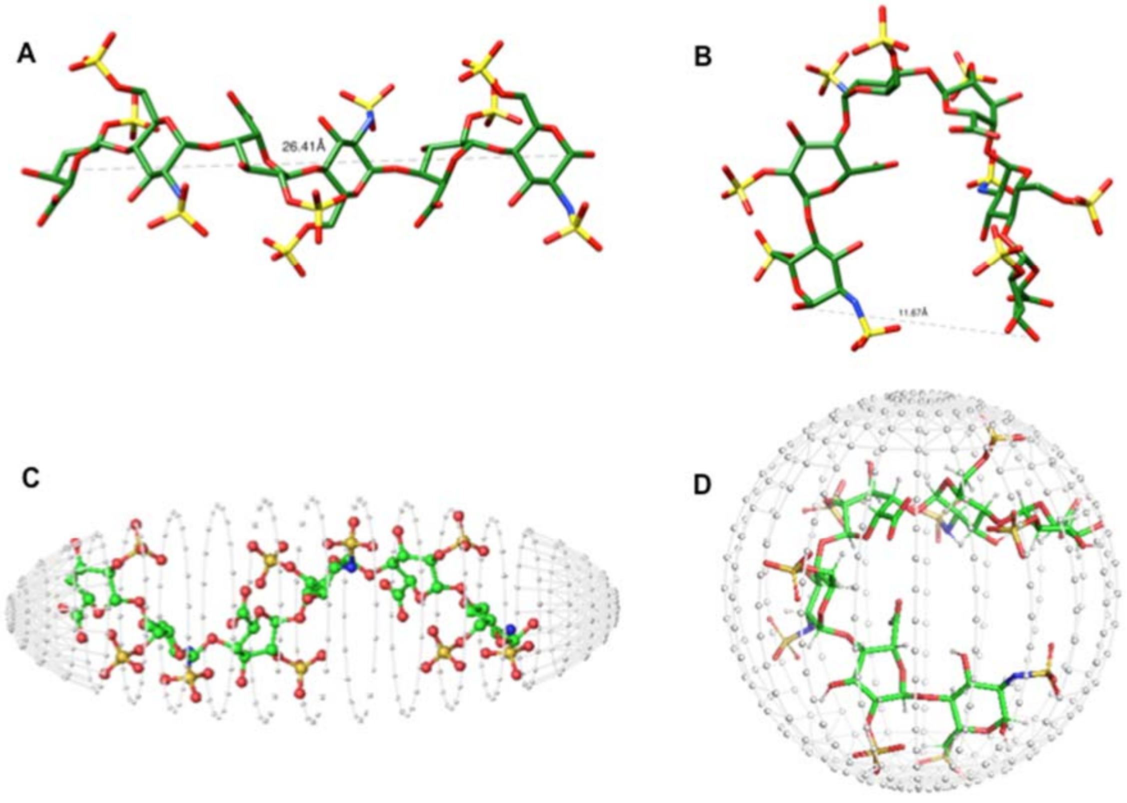


Figure S2. A) The value of end-to-end distance EED for a HS06 rod like structure shown from PDBID:1HPN. B) The value of end-to-end distance EED for a theoretically possible HS06 hairpin bent structure. C) Shows the MVEE enclosing HS06 rod like structure, D) Enclosing MVEE shown for a HS06 in hairpin bent structure (in both C and D HS06 shown in green color with enclosing ellipsoid grid points in grey color).

| #A | AVERAGE VALUES (Å) | | | |
|---|---|---|---|---|
| | GLYCAM06 | CHARMM36 | NMR Hexasaccharide (PDBID :1HPN) | Hairpin structure |
| EED | 26.52 | 25.50 | 26.4 | 11.67 |

| #B | AVERAGE VALUES ($Å^3$) | | | |
|---|---|---|---|---|
| | GLYCAM06 | CHARMM36 | NMR Hexasaccharide (PDBID :1HPN) | Hairpin structure |
| MVEE | 773.32 | 797.26 | 673.90 | 643.38 |
| R1 | 6.37 | 6.39 | 6.120 | 6.41 |
| R2 | 7.46 | 7.41 | 7.29 | 9.90 |
| R3 | 16.33 | 16.88 | 15.11 | 10.14 |

Table S1. A) Average value of end-to-end distance EED and B) average value of minimum volume enclosing ellipsoid MVEE from GLYCAM06 and CHARMM36 force fields for the HS06 dynamics in explicit water. For comparison the values of HS06 from 1HPN structure and theoretically built hairpin structures are shown.

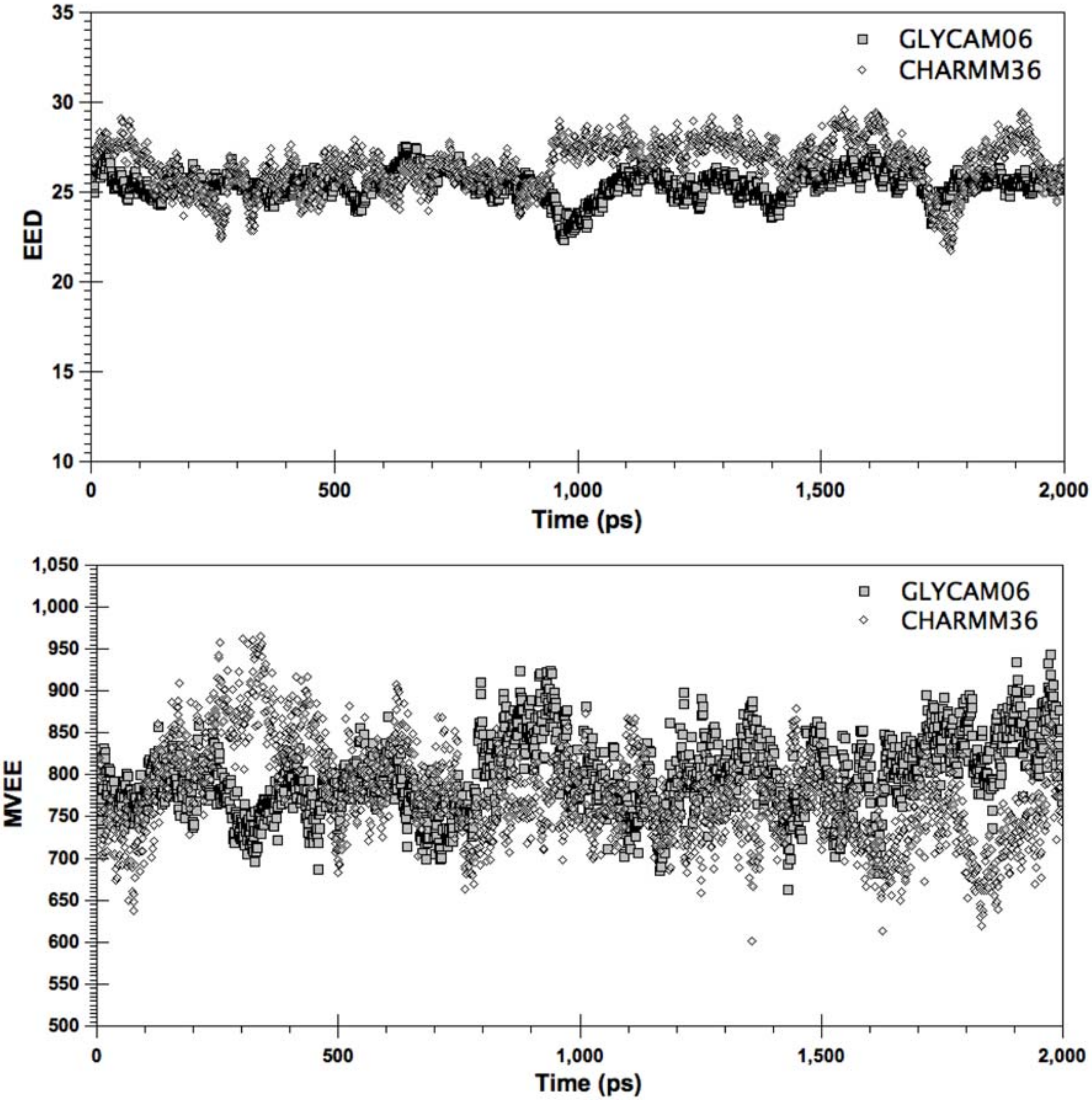


Figure S3. End-to-end distance EED (top) and Minimum volume enclosing ellipsoid MVEE (bottom) from GLYCAM06 and CHARMM36 force fields for the HS06 dynamics in explicit water with respect to time frame in each ps from last 2ns. Typically showing overlap and non overlap regions with definite range.

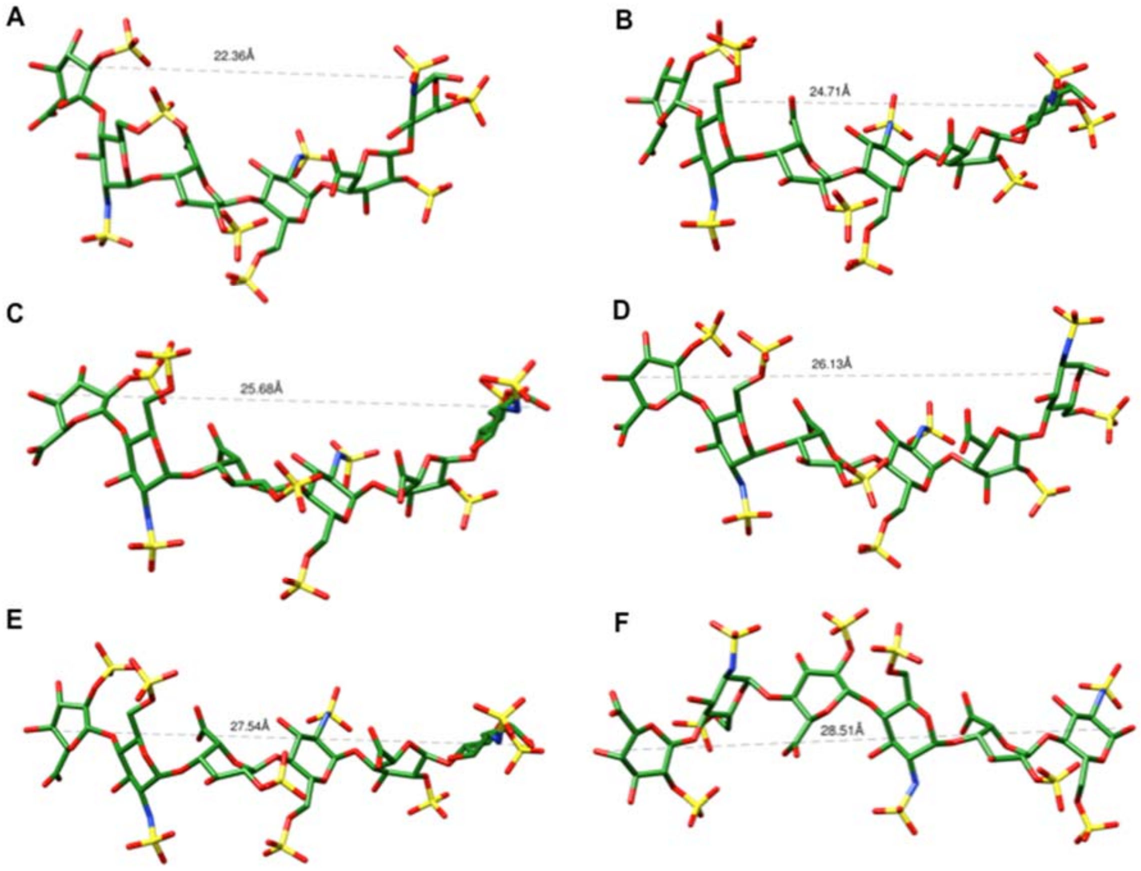


Figure S4. A) to F) shows the representative conformers from the bent to more rigid road like structure of HS06 based on the end-to-end distance (EED) From the MD generated conformers.

**IdoA2S-GlcNS6S**

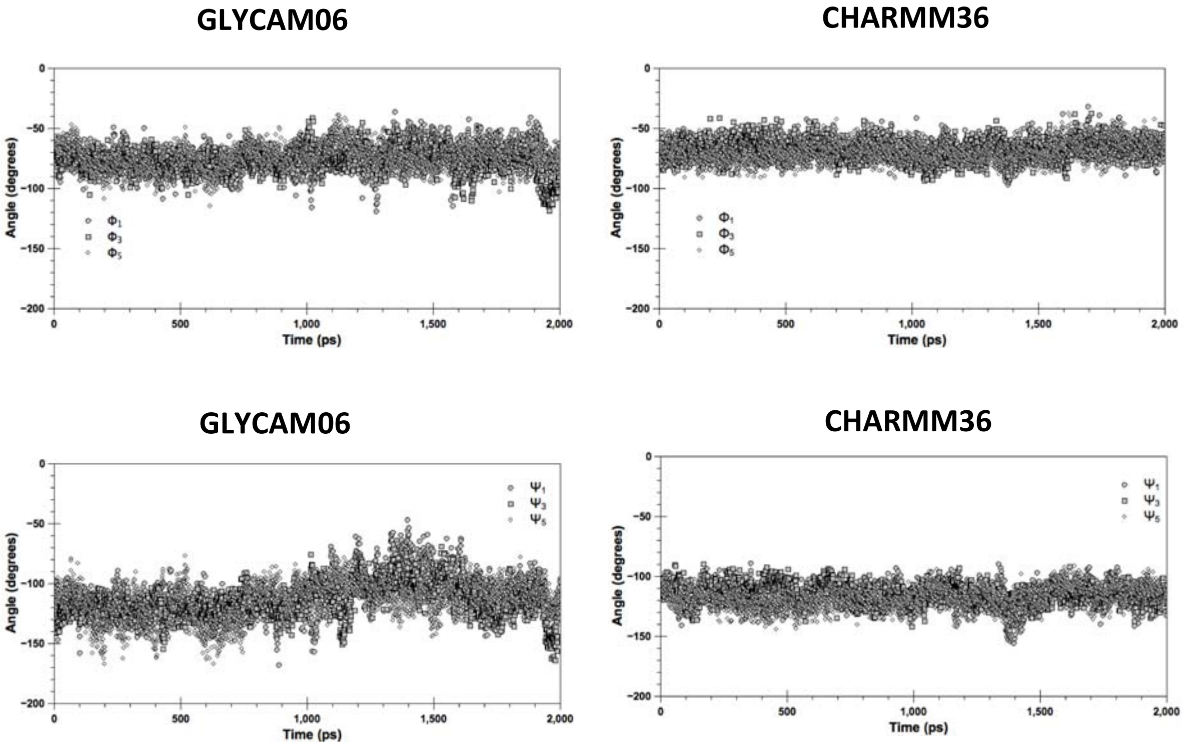


Figure S5. Phi and psi values for the last 2ns from the glycosidic linkage of IdoA2S-GlcNA6S from GLYCAM06 and CHARMM36 force fields for the HS06 dynamics in explicit water with respect to time frame 1ps

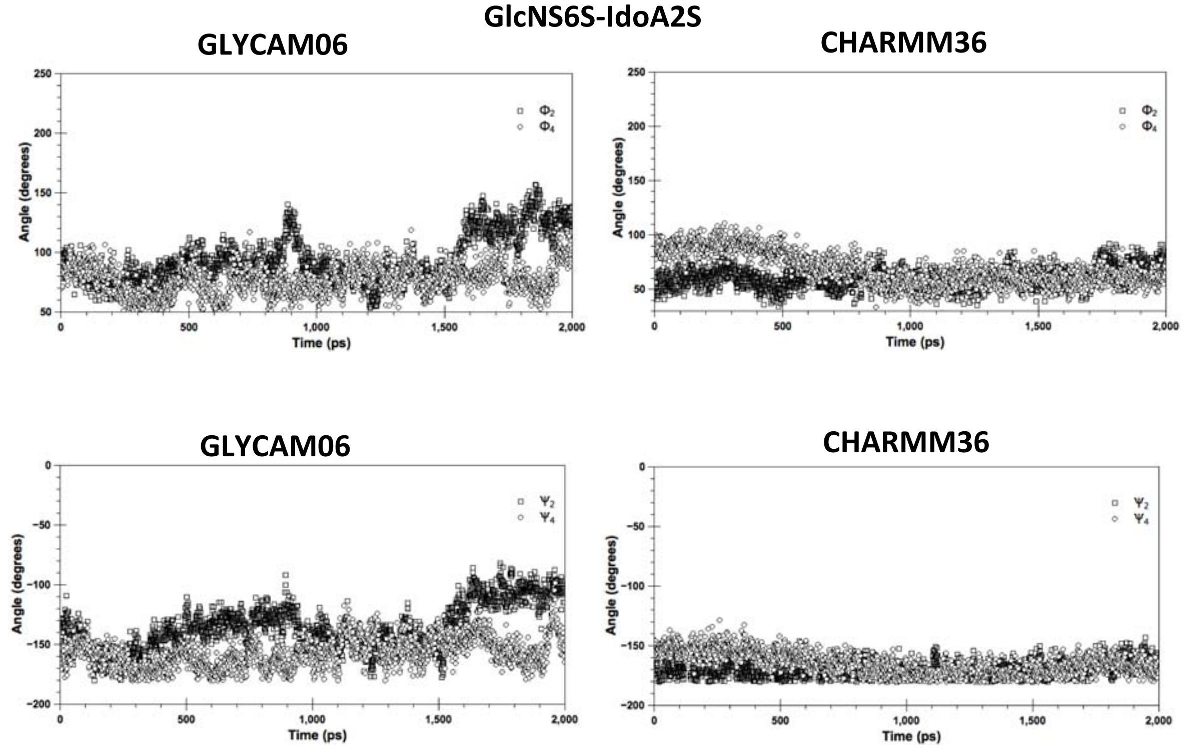


Figure S6. Phi and psi values for the last 2ns from the glycosidic linkage of GlcNA6S-IdoA2S rom GLYCAM06 and CHARMM36 force fields for the HS06 dynamics in explicit water with respect to time frame 1ps

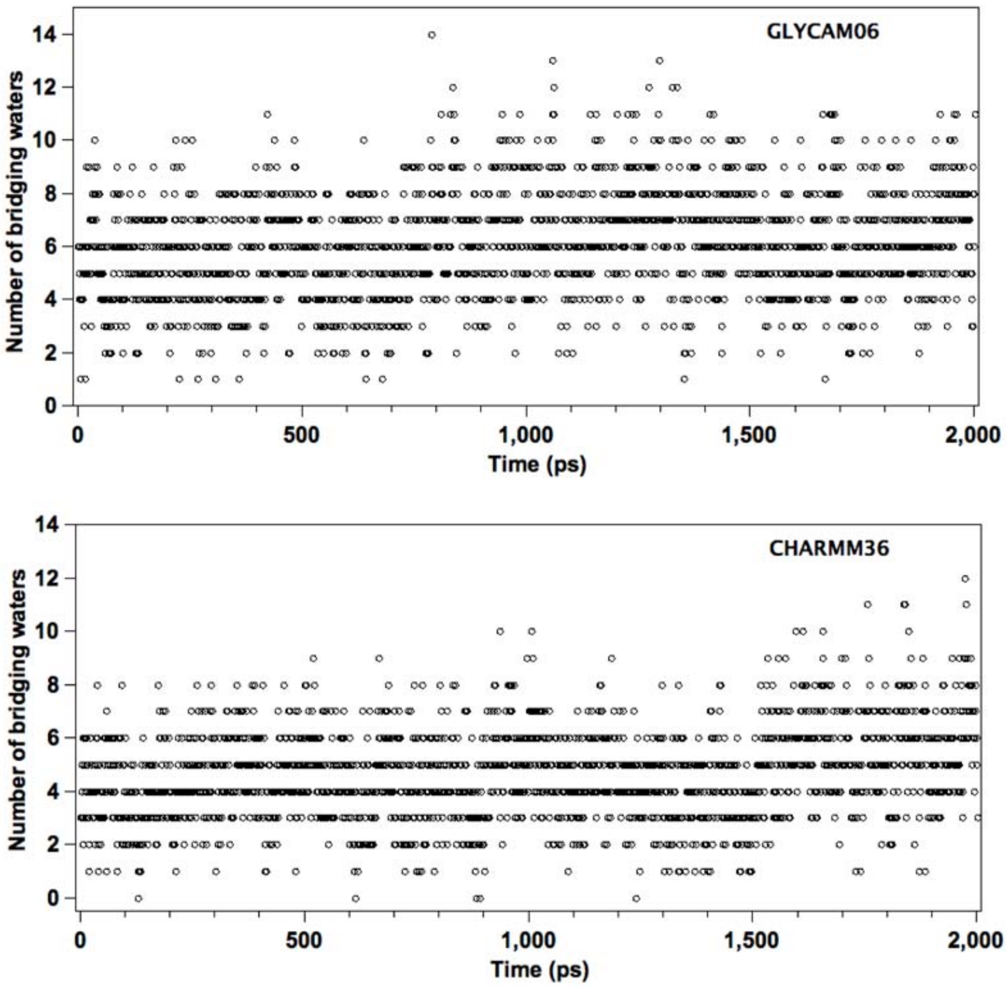


Figure S7. The numbers of bridging water molecules between the acceptor and donors of HS06 residue for each time frame are shown for GLYCAM06 and CHARMM36 force fields.
